# Control of lineage-specific gene expression by functionalized gRNA barcodes

**DOI:** 10.1101/178897

**Authors:** Aziz Al’Khafaji, Amy Brock

## Abstract

Lineage tracking delivers essential quantitative insight into dynamic, probabilistic cellular processes, such as somatic tumor evolution and differentiation. Methods for high diversity lineage quantitation rely on sequencing the population of DNA barcodes. However, molecular analysis of specific individual lineages is not possible with this approach. To address this challenge, we developed a functionalized lineage tracing tool that enables high diversity lineage tracing and lineage-specific manipulation of gene expression. This modular platform utilizes expressed barcode gRNAs to both track cell lineages and direct lineage specific gene expression.

## Introduction

Many pathological and physiological processes, including cancer, infection, and microbiota control, are governed by the evolutionary dynamics of large heterogeneous cell populations. Tumors consist of 10^7^-10^12^ cells that vary with respect to growth rate, drug response, and cell fate decisions. While rare mutations are a driving force for population adaptation, new evidence also emphasizes the contribution of epigenetic plasticity and heterogeneous cell states within clonal populations. Intratumor cell heterogeneity is a significant clinical challenge that contributes to chemoresistance and treatment failure^1-10^. To improve therapeutic strategies in cancer and infectious diseases, new tools are required to measure and control the contributions of heterogeneous cell populations ^11-14^.

Recent studies have demonstrated the utility of high-diversity DNA barcode libraries in monitoring heterogeneous cell populations ^11-18^. This is achieved by labeling each cell in a population with a unique, random, heritable sequence; lineage abundance is tracked over time by next-generation sequencing of the barcode ensemble. Changes in clonal dynamics after perturbations, such as treatment with a pharmacological agent, may reveal variation in lineage survival or proliferation rate^13-16^. This approach allows for the simultaneous observation of many cell lineage trajectories to reveal high-resolution details of population dynamics^18^. However, quantitation of lineage abundance by sequencing is a destructive measurement that limits further molecular and functional analysis of the cells in specific lineages of interest. We are not aware of any existing method that enables one to simultaneously track high diversity lineage dynamics and isolate a specific lineage from a cell population for downstream analysis. Here we set out to develop a platform technology, **C**ontrol **o**f **L**ineages by **B**arcode **E**nabled **R**ecombinant **T**ranscription (COLBERT), to enable lineage tracing followed by lineage-specific activation of gene expression.

In this technique a population of cells are tagged with a stably integrated barcode-gRNA under control of a constitutive promoter. Lineage-specific gene expression can then be accomplished by transfecting the entire population of cells with a plasmid containing a transcriptional activator variant of Cas9, dCas9-VPR, and a “Recall” plasmid encoding the lineage barcode of interest upstream of a gene to be activated ^19^. Only those cells containing the specified barcode-gRNA of interest, in coordination with dCas9-VPR, drive expression of the reporter gene. A schematic of the overall strategy of COLBERT is shown in **Fig. 1a**.

**Figure 1.**
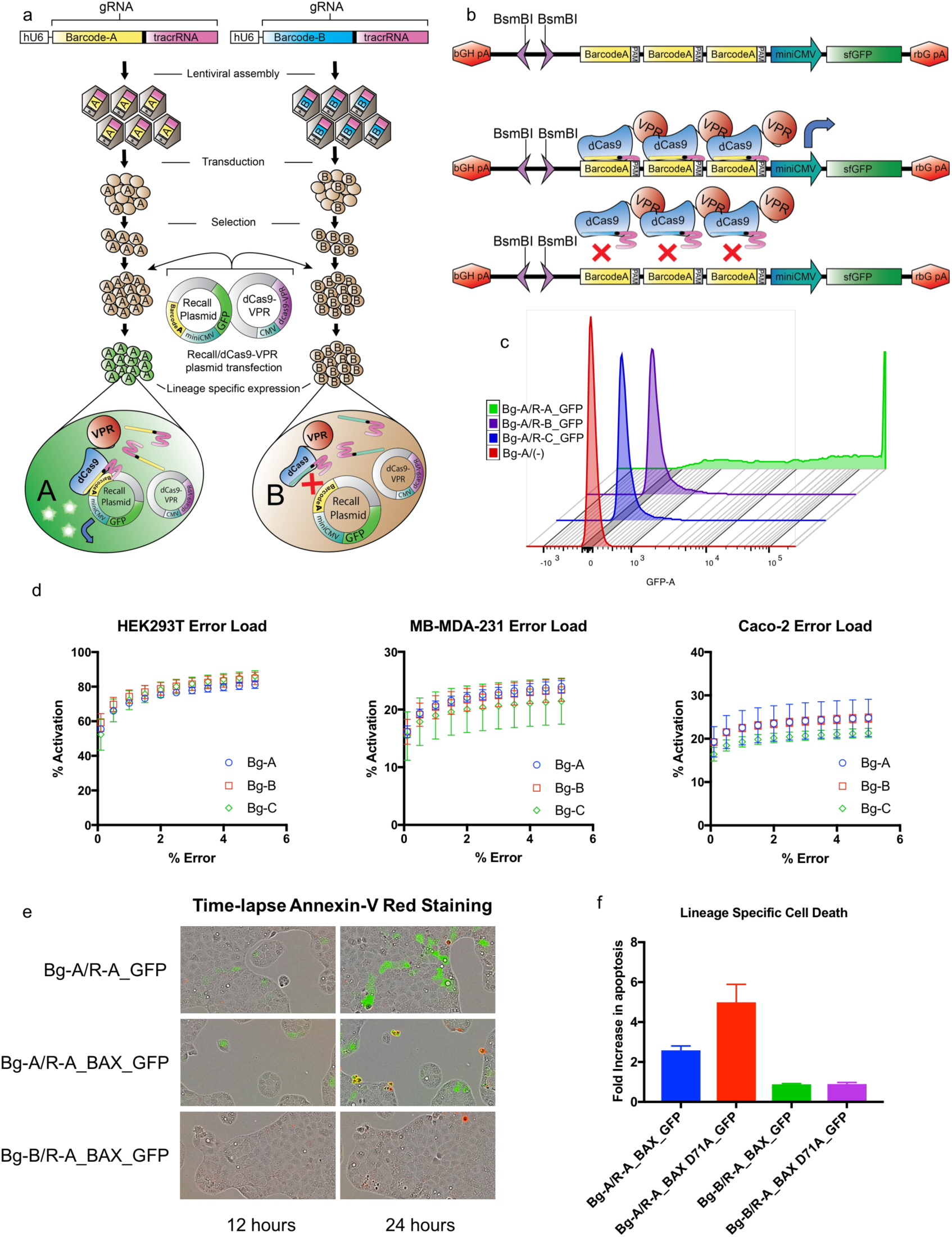
Lineage-specific activation of gene expression. (a) Generation and lineage specific gene activation of independent barcoded gRNA populations. Three unique barcodes were randomly generated following the GNSNWNSNWNSNWNSNWNSN template and assembled into lentiviral gRNA expression cassettes. Cell lines: HEK293T, Caco2, and MDA-MB-231 were independently transduced with the three different barcode gRNAs and selected for stable integration. The barcoded populations were then co-transfected with each of one of the Recall plasmids, R-A/B/C_GFP and the dCas9-VPR plasmid. GFP expression was assessed 48 h post transfection via flow cytometry. (b) View of the lineage specific expression components. The lineage-specific Recall Plasmid contains a 3x barcode of interest_PAM array and adjacent downstream miniCMV promoter_sfGFP. In the presence of the matching barcode gRNA/dCas9-VPR complex, binding of the barcode arrays by the transcriptional activator dCas9-VPR will drive expression of sfGFP. In the case of mismatching barcode gRNA/dCas9-VPR complex, binding of the barcode arrays will not occur and expression of sfGFP will not be driven. (c) staggered histograms comparing high GFP expression for instances of matching barcode gRNA/Recall plasmid and nominal expression for instances of mismatch. GFP expression was measured via flow cytometry. (d) Error load graphs showing percent positive population activation at a given error rate. (e) Time-lapse fluorescent imaging of Caco2 Bg-A and Bg-B populations transfected with Recall-A_GFP and Recall-A_BAX_GFP. The matching Bg-A population transfected with R-A_BAX_GFP displays efficient gene expression with GFP expression starting at 12 h followed BAX induced apoptosis present at 24 h. The mismatched Bg-B population does not show either GFP expression or notable induction of apoptosis. (f) Quantification of Annexin-V red shows that, relative to Bg-A/R-A_GFP, there is a significant fold increase apoptosis in the cell populations transfected with a matching Recall plasmid containing either BAX or the hyper active mutant BAX D71A. Additionally, neither population with a mismatching Recall plasmid containing BAX or the hyperactive mutant displayed an increase in apoptosis over the control.

## Results

To demonstrate lineage-specific expression of a fluorescent reporter by COLBERT, we generated three independent HEK293T cell populations expressing a single known barcode gRNA (Bg), Bg-A, Bg-B, Bg-C. Cells were transduced at a multiplicity of infection (MOI) of 0.1 to minimize instances of integration of more than one barcode. Cells containing stably integrated barcode sequences were selected by BFP^+^ expression (Fig. 1a). We then constructed three different Recall plasmids (Recall A-C) each containing one of the three corresponding barcode regions and PAM site upstream of a miniCMV promoter and sfGFP (**Fig. 1b**). We then co-transfected these barcode populations with each of the Recall plasmids + dCas9-VPR plasmid independently, causing instances of either match or mismatch with regards to the barcoded gRNA and Recall plasmid (**Fig. 1a**). After 48 h, GFP expression was assessed via flow cytometry. Our results showed that barcoded cell populations transfected with a matching Recall plasmid were able to activate expression of the fluorescent reporter, while only nominal expression was present in the instances of mismatch (**Fig. 1c**). This robust and easy-to-use platform can be deployed in a variety of cell types. To assess the efficiency of lineage-specific GFP activation in the match population and compare with non-specific activation of mismatch population, we quantified the error load associated with deploying the system in HEK293T, Caco2, and MB-MDA-231 cells. 75% of the lineage-specific GFP^+^ cells could be identified in HEK293T with 2% false positive activation. Error rates also remained low in Caco2, and MB-MDA-231 cells, although GFP activation was significantly lower, predominantly due to less efficient plasmid transfection in these cell types (**Fig. 1d**). Additionally, discrepancies in transfection efficiency and lineage specific gene activation could be due to the efficiencies of the dCas9-activator of choice in the specific cell type ^20^.

To optimize lineage-specific activation with the Recall plasmid, alternative designs were tested with varying numbers of barcode recognition sites (1x, 3x and 6x). In addition, both Recall and dCas9 VPR plasmid were titrated to determine optimal amounts to activate barcode-driven expression. (**Suppl Fig S1-S3**)

Beyond the control of fluorescent reporter gene expression, this system can be functionalized to express any set of genes in a lineage-specific manner. To explore the multifunctionality of this platform we sought to perturb the cell fates of specific lineages, by driving lineage-specific expression of the pro-apoptotic protein BAX and the hyperactive mutant BAX D71A, in conjunction with GFP ^21^. Time-lapse fluorescent imaging reveals lineage-specific gene expression of GFP and subsequent apoptosis of fluorescing cells (**Fig. 1e**). Co-staining for annexin confirms activation of apoptotic signaling, with increased levels of apoptosis occurring in the BAX D71A mutant (**Fig. 1f)**.

To confirm the specificity and efficiency of lineage-specific expression, we tested the tool in a large diverse barcoded population. We constructed a high-diversity barcode gRNA library with the template: GNSNWNSNWNSNWNSNWNSN, having a diversity potential greater than 500,000,000 unique sequences (**Fig. 2a**). This gRNA library was ligated into a gRNA expression lentiviral transfer vector and assembled into a pooled gRNA barcoded lentivirus. Following transduction, stably integrated BFP^+^ cells were collected, yielding a high diversity population of <10^6^ barcoded cells. Cells from the Bg-A population were then spiked into the high diversity library at 1:100 and 1:1000 dilution and grown overnight. The spiked populations were then co-transfected with the Recall-A_GFP and dCas9-VPR plasmids and sorted via FACS for GFP expression. Sorted cells were subcultured and genomic DNA was isolated for sequencing (**Fig. 2a**). Barcode sequences were amplified using PCR and analyzed by Sanger sequencing. Barcode sequencing of the population confirms that COLBERT identified and allowed for the isolation of the fraction of cells carrying the reference Bg-A barcode from within the high diversity Bg-Library population (**Fig. 2b)**.

**Figure 2.**
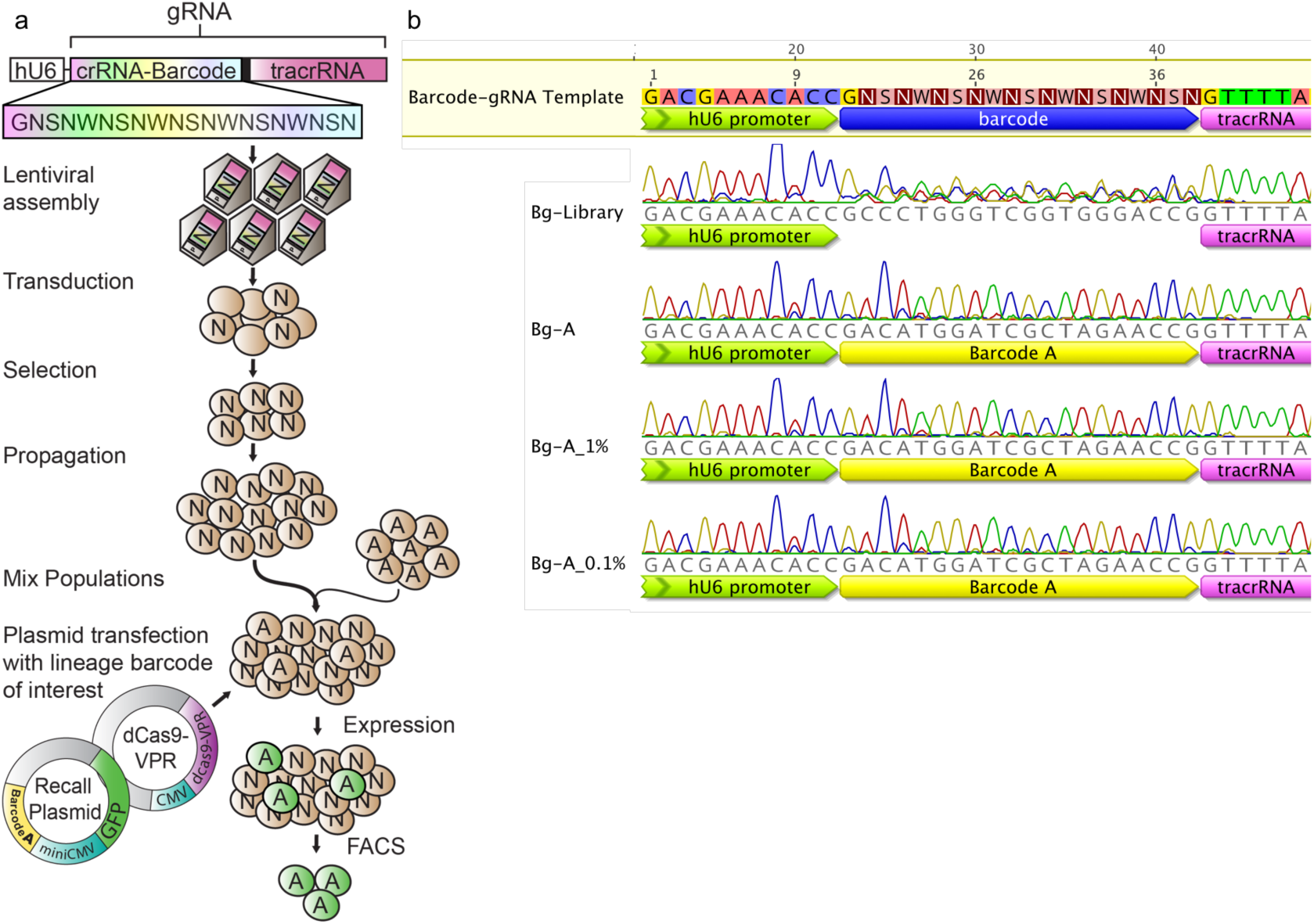
Isolation and manipulation of a single lineage of interest within a high diversity population. High diversity barcoded-gRNA HEK293T cell population was generated with a GNSNWNSNWNSNWNSNWNSN template. The HEK293T Bg-A population was spiked in with the high diversity population to obtain a 1% and 0.1% Bg-A mixed population. Bg-A cells were then isolated from the mixed population via co-transfection of the R-A_GFP plasmid and dCas9-VPR plasmid and cell sorted based off of GFP expression. (b) Sequencing data of the Bg-library and Bg-A populations as well as the Bg-A 1% and 0.1% isolated populations. The Bg-A 1% and 0.1% isolated population sequencing shows a near identical chromatogram sequence to the parent 100%Bg-A population, displaying the capacity of COLBERT to identify lineages of interest for a highly complex background.

The demonstration that expressed gRNA barcodes can be used to efficiently perform lineage-specific manipulation of gene expression opens up the possibility for a broad range of studies investigating the potential of lineage-specific perturbations within the context of a heterogeneous, evolving cell population (**Fig. 3)**. Following barcode instantiation, cells are permitted to proliferate and at intervals the genomically encoded barcode region may be sequenced for quantitation of clonal barcodes; a parallel sample portion can be archived for retroactive analysis. Lineage dynamics may inform the identification of specific lineages of interest for subsequent gene activation in archival samples. The ability to concurrently track clonal fitness dynamics and generate lineage-specific genomic and transcriptomic data over longitudinal studies will provide unprecedented insight into cancer adaptation and other diseases with an evolutionary basis.

**Figure 3.**
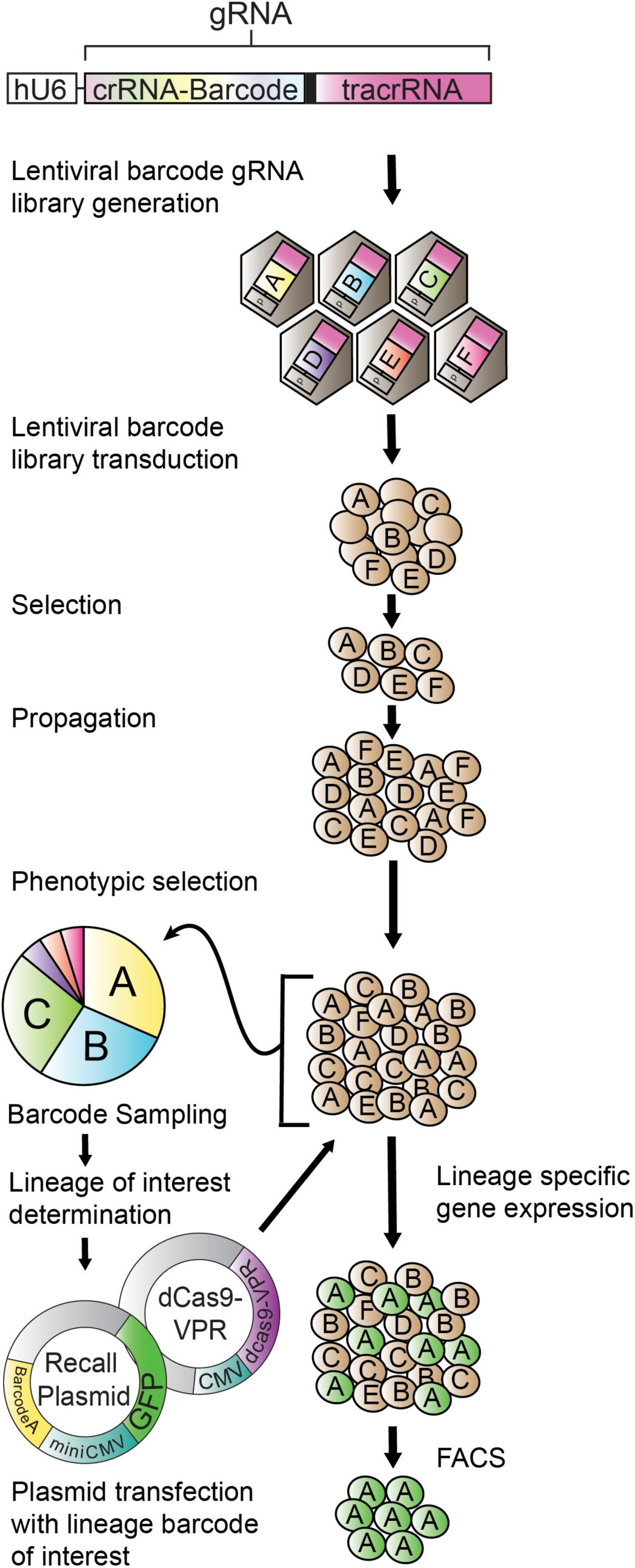
Example experimental workflow utilizing COLBERT. A population of cells are tagged with a library of expressed barcoded gRNAs. The barcoded population can subsequently go through a phenotypic selection of interest. The resulting selected population is then sampled for relative barcode abundance. The barcode analysis would inform lineages of interest and allow for lineage-specific Recall plasmids to be assembled and co-transfected with dCas9-VPR into the resultant population for lineage-specific gene expression. In this instance, GFP is expressed, allowing the lineage of interest to be cell sorted and isolated from the mixed population.

## Methods

### High-complexity Barcode-gRNA Library Construction

The following 60 base-pair oligonucleotide containing a 19 nucleotide semi-random sequence corresponding the barcode guide-RNA and reverse extension primer was ordered from Integrated DNA Technologies:

GAGCCTGAAGACCTCACCGNSNWNSNWNSNWNSNWNSNGTTTTAGCGTCTT CCATGCGCA, TGCGCATGGAAGACGCTAAAAC. An extension reaction was performed to generate the double stranded barcode-gRNA oligo. The double stranded product contains two BbsI sites that, after digestion, generate complimentary overhangs for ligation into the gRNA expression transfer vector pKLV-U6gRNA(BbsI)-PGKpuro2ABFP (Addgene). 1μg of BbsI digested gRNA expression transfer vector was ligated with digested barcode-gRNA insert in a molar ratio 1:7. This reaction was cleaned and concentrated in 6μl using the Zymo DNA Clean & Concentrator™ kit and transformed into electrocompetent SURE 2 cels (Agilent). Transformants were inoculated into 500ml of 2xYT containing 100μg/ml carbenicillin for outgrowth overnight at 37 °C. Transformation efficiency was calculated via dilution plating and shown to be approximately 7 x 10^8^ cfu/μg.

### Mock Barcode-gRNA construction

Three different discrete known barcode-gRNA lentiviral expression vectors were generated with the sequences: A) GACATGGATCGCTAGAACCG, B) GTCAAGGTAGCTAAGTAGCG, C) GTCAAGCGTGCAATGGTAGC. To accomplish this, oligo pairs with complimentary barcode sequences and the appropriate overhang sequences were mixed and cloned into the BbsI digested pKLV-U6gRNA(BbsI)-PGKpuro2ABFP transfer vector at a 10:1 molar ratio:

A) CACC**GACATGGATCGCTAGAACCG**GT,

TAAAAC**CGGTTCTAGCGATCCATGTC**,

B) CACC**GTCAAGGTAGCTAAGTAGCG**GT,

TAAAAC**CGCTACTTAGCTACCTTGAC**,

C) CACC**GTCAAGCGTGCAATGGTAGC**GT,

TAAAAC**GCTACCATTGCACGCTTGAC**.

### Lentiviral Assembly

Lentiviral assembly was accomplished using the GeneCopeia Lenti-Pac™ HIV Expression Packaging Kit (cat# HPK-LvTR-20). Two days prior to lentiviral transfection HEK293T cells were plated onto a 10cm dish at 1.5 million cells and cultured in 10ml DMEM supplemented with 10% heat inactivated FBS. 48 h after plating, cells were 70-80% confluent and transfected with 15μl of EndoFectin and a mix of 2.5μg of pKLV-U6**Barcode-gRNA**-PGKpuro2ABFP and 2.5μg of Lenti-Pac™ HIV mix (GeneCopoeia). The media was replaced 14 h post transfection with 10ml DMEM supplemented with 5% heat inactivated FBS and 20μl TiterBoost™(GeneCopoeia) reagent. Media containing viral particles was collected at 48 and 72 h post transfection, centrifuged at 500g for 5 min, and filtered through a 45μm polyethersulfone (PES) low protein-binding filter. Filtered supernatant was aliquoted and stored at −80°C for later use.

### Barcoding cell lines

Cell lines HEK293T, MB-MDA-231, and Caco-2 cell lines were cultured in DMEM medium supplemented with 10% FBS and 1% penicillin-streptomycin. Cells were transduced with the pKLV-U6**Barcode-gRNA**-PGKpuro2ABFP lentivirus using 1μg/ml polybrene. After 48 h incubation, BFP^+^ cells were isolated by FACS. To reduce the likelihood that two viral particles enter a single cell, the lentiviral transduction multiplicity of infection was kept below 0.1.

### Barcode amplification

After lineage isolation, cell populations of interest were harvested and genomic DNA was extracted using the PureLink^®^ Genomic DNA Mini Kit (Thermo Fisher). Barcode sequences were amplified using PCR and sent for Sanger sequencing. For each PCR reaction, 250ng of genomic DNA was used as a template.

### Recall Plasmid Assembly

The Recall plasmid was constructed by using standard restriction cloning to combine a gBlock^®^ containing three tandem type IIS restriction sites (BsmBI, BbsI, BsaI) flanked by terminators with an amplicon containing a bacterial replication origin and ampicillin resistance marker to create this Golden Gate ready vector. Genes and barcode-specific landing pad sequences were cloned into the recall plasmid using the type IIS restriction sites. Barcode-specific landing pad arrays were generated by ordering phosphorylated complimentary oligo pairs, corresponding with the barcode sequence of interest, with specific overlaps that both direct assembly of the landing pad array and integration into the Recall plasmid (**Suppl Fig S1**). The landing pad arrays were ligated and gel extracted to ensure cloning with a fully assembled array. The fully assembled barcode landing pad was cloned into the BbsI site using standard restriction digest cloning. Mock Recall screens were used to assess efficiency via lineage specific expression of sfGFP. This reporter construct was assembled by cloning in a gBlock^®^ encoding miniCMV-sfGFP into the BsaI site using Golden Gate Assembly (Recall_GFP). Lineage-specific cell death was measured via barcode driven expression BAX and the hyper active mutant BAX D71A. gBlocks^®^ encoding miniCMV-BAX and miniCMV-BAX D71A were cloned into the BsmBI sites using Golden Gate Assembly (Recall_BAX_GFP, Recall_BAX D71A_GFP).

### Mock Recall Screens

The mock screens were performed in 24 well plates. HEK293T cells were transfected at 60% confluence using 1.5μl Lipofectamine™3000, 1μl P3000™ Reagent, 150ng of Recall_GFP plasmid and 500ng of dCas9-VPR plasmid. Caco2 cells were transfected at 30% confluence and transfected using 1μl Lipofectamine™LTX, 0.5μl Plus™ Reagent, 250ng Recall_GFP Plasmid, and 250ng dCas9-VPR plasmid. MB-MDA-231 cells were transfected at 70% confluence using 1μl Lipofectamine™LTX, 0.5μl Plus™ Reagent, 250ng Recall_GFP Plasmid, and 250ng dCas9-VPR plasmid. Cells were analyzed for GFP expression via flow cytometry 48 h post-transfection. Error load was quantified by comparatively tallying the values of the GFP-histogram, from high to low GFP intensity, of matching and mismatching recall samples. Specifically, sum totals of matching recall events were tabulated with respect to and along with each new accruing mismatch recall event. From these tabulations, % activation at a given error was calculated using the formula:

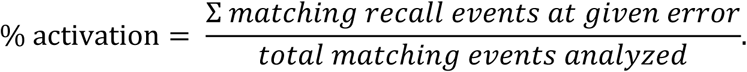

% error was calculated using the formula:

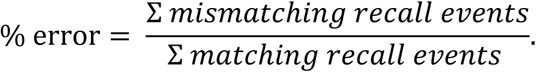

The calculated % activation and % error values were plotted on a scatter plot plot and, using least squares fitting, generated exponential equations to model the data. (**Suppl. Fig. S4**). Fits for a vast majority of samples had an R^2^ value of > 0.95. Experiments were performed in biological triplicate and modeled error load values were averaged along with standard error calculated.

### Lineage Isolation

For a standard, a range of HEK293T Bg-A/Bg-library dilutions were plated in 35 mm tissue culture dishes (100%Bg-A, 50% Bg-A, 10% Bg-A, 1% Bg-A, 0.1% Bg-A) with total cell number 360,000 per well. Two 10cm plates were plated at 2.2 million cells for both a 1% and 0.1% Bg-A lineage dilution for lineage isolation. The 35 mm dishes were transfected with 4.5μl Lipofectamine™LTX, 2.25μl Plus™ Reagent, 675ng Recall Plasmid, and 1.575μg dCas9-VPR plasmid per well. The 10cm plates were transfected with 27.5μl Lipofectamine™LTX, 13.75μl Plus™ Reagent, 4.125μg Recall Plasmid, and 9.625μg dCas9-VPR plasmid per plate. Populations were flow analyzed and sorting gates were set using 0% Bg-A as a standard (**Suppl Fig. S5**). The GFP^+^ population was cultured for isolation of genomic DNA as above.

### Annexin V Red Assay

Caco2 were transfected at 30% confluence using 1μl Lipofectamine™LTX, 0.5μl Plus™ Reagent, 250ng Recall Plasmid, and 250ng dCas9-VPR plasmid. At time of transfection, 2.5μl IncuCyte^®^ Annexin V Red Reagent (Essen BioScience Cat # 4641) was added to monitor apoptosis. Cells were monitored in the IncuCyte^®^ for real time measurement of apoptotic cells in culture via fluorescent quantitation. Images were collected every 2 h and quantitation of apoptotic was performed using the IncuCyte^®^ image analysis software.

## Acknowledgements

This work was supported by a Texas 4000 Foundation Cancer Research Pilot Grant (to A.B.). The authors are grateful to Amy Price for excellent administrative support.

## References

1. Greaves, M. Evolutionary determinants of cancer. Cancer Discov. 5, 806–821 (2015).

2. Brock, A., Chang, H. & Huang, S. Non-genetic heterogeneity--a mutation-independent driving force for the somatic evolution of tumours. Nat Rev Genet 10, 336-342 (2009).

3. Sharma, S. V. et al. A chromatin-mediated reversible drug-tolerant state in cancer cell subpopulations. Cell 141, 69-80 (2010).

4. Huang, S. & Kauffman, S. How to escape the cancer attractor: rationale and limitations of multi-target drugs. Semin Cancer Biol 23, 270-278 (2013).

5. Polyak, K. Tumor Heterogeneity Confounds and Illuminates: A case for Darwinian tumor evolution. Nat Med 20, 344-346 (2014).

6. Archetti, M., Ferraro, D. A. & Christofori, G. Heterogeneity for IGF-II production maintained by public goods dynamics in neuroendocrine pancreatic cancer. Proc Natl Acad Sci U S A 112, 1833-1838 (2015).

7. Grosse-Wilde, A. et al. Stemness of the hybrid Epithelial/Mesenchymal State in Breast Cancer and Its Association with Poor Survival. PLoS One 10, e0126522 (2015).

8. Cleary, A. S., Leonard, T. L., Gestl, S. A. & Gunther, E. J. Tumour cell heterogeneity maintained by cooperating subclones in Wnt-driven mammary cancers. Nature 508, 113-117 (2014).

9. Quintana, E. et al. Phenotypic heterogeneity among tumorigenic melanoma cells from patients that is reversible and not hierarchically organized. Cancer Cell 18, 510-523 (2010).

10. Pisco, A. O. et al. Non-Darwinian dynamics in therapy-induced cancer drug resistance. Nat Commun 4, 1–11 (2013).

11. Omics, S. Building a lineage from single cells: genetic techniques for cell lineage tracking. Nat Rev Genet. (2017).

12. Frieda, K. L. et al. Synthetic recording and in situ readout of lineage information in single cells. Nature, 541, 107–111 (2017).

13. Rogers, Z. N. et al. A quantitative and multiplexed approach to uncover the fitness landscape of tumor suppression in vivo. Nat Met. 14, (2017).

14. Yu, C. et al. High-throughput identification of genotype-specific cancer vulnerabilities in mixtures of barcoded tumor cell lines. Nat Biotechnol 34, (2016).

15. Bhang H. E. et al. Studying clonal dynamics in response to cancer therapy using high-complexity barcoding. Nat Med, 21, 440-8 (2015).

16. Hata A. N. et al. Tumor cells can follow distinct evolutionary paths to become resistant to epidermal growth factor receptor inhibition. Nat Med. 22, 262-269. (2016)

17. Levy, S. F. et al. Quantitative evolutionary dynamics using high-resolution lineage tracking. Nature 519, 181–6 (2015).

18. Blundell, J. R. & Levy, S. F. Genomics Beyond genome sequencing: Lineage tracking with barcodes to study the dynamics of evolution, infection,and cancer. Genomics 104, 417–430 (2014).

19. Chavez, A. et al. Highly efficient Cas9-mediated transcriptional programming. Nat Meth 12, 326–328 (2015).

20. Chavez, A. et al. Comparison of Cas9 activators in multiple species. Nat Met 13, 7-10 (2016).

21. Zhou, H., Hou, Q., Hansen, J. L. & Hsu, Y. Complete activation of Bax by a single site mutation. Oncogene 26, 7092–7102 (2007).

